# Distinct neural correlates for focusing on similar memory contents originating from current or previous experience

**DOI:** 10.64898/2026.06.05.730490

**Authors:** Dongyu Gong, Dejan Draschkow, Anna C. Nobre

## Abstract

Flexible, goal-directed behavior depends on the ability to select and prioritize information from memory representations freshly encoded from the sensory stream as well as retrieved from previous experience. The spatial gating signatures of internal attention in working memory (WM) are increasingly well characterized, but it remains unclear whether the same neurophysiological mechanisms are recruited for orienting attention to items from long-term memory (LTM). We recorded EEG and eye movements while participants focused on WM versus LTM representations in a unified precision-report task. Retrocues improved performance for both WM and LTM items, but with larger behavioral gains for WM items. Neural and oculomotor markers of spatial orienting, including contralateral posterior alpha suppression and gaze biases toward remembered locations, were robust when focusing on WM items; but were reliably weakened or absent when focusing on LTM items. Multivariate pattern analyses provided complementary evidence for the recruitment of dissociable neural mechanisms when focusing on WM vs. LTM items, which unfolded with similar time courses. Together, the results establish the existence of dissociable neural mechanisms for internal attention, which can be deployed flexibly depending on the relevant memory trace to guide performance. The findings raise interesting and tractable questions about whether differences in representational formats or representational domains drive the distinct internal attention mechanisms.

## Introduction

The remarkable capacity of the human mind to guide behavior is fundamentally reliant on memory (Hutchinson & Turk-Browne, 2012; Nobre & Stokes, 2019; Schacter et al., 2007). To navigate complex and dynamic environments, we constantly draw upon our internal store of knowledge, selectively accessing information that is relevant to our current goals. Selectively focusing within internal mental representations is termed internal attention, contrasting with external attention, which operates on incoming information from the sensory stream (Chun et al., 2011).

Most of our knowledge about internal attention comes from investigating how we select and prioritize contents encoded into working memory from the sensory stream (see Chun et al., 2011; Souza & Oberauer, 2016; Van Ede & Nobre, 2023). How internal attention operates on contents stored within long-term memory to guide selective retrieval is less well understood (see Cabeza et al., 2008; Chun et al., 2011; Long & Kuhl, 2019). For example, orienting attention to contents in visual working memory encoded from the sensory stream consistently relies on spatial information, even when the task is entirely non-spatial in nature and does not strictly require reporting of any spatial attributes (Pertzov & Husain, 2014; Schneegans et al., 2021). Neurophysiology studies show spatially specific modulation of alpha-band activity (e.g., van Ede et al., 2017), and oculomotor studies show systematic spatial biasing of eye gaze and microsaccades (van Ede, Chekroud, & Nobre, 2019). Whether similarly spatial anchoring occurs when retrieving non-spatial attributes from long-term memory is unknown.

In a recent study, we developed a task design to compare the behavioral consequences of internal selective attention for selecting contents encoded from the sensory stream vs. retrieved from long-term memory within the same trials (Gong et al. 2025). Participants learned the colors and shapes of items at two locations. In a subsequent attention-orienting task, two other colored shapes appeared on trials and were encoded into working memory. At the end of the trial, participants had to reproduce the shape of one of the items from long-term memory or from the sensory stream. Informative colored retrocues during the delay indicated the item to be reported, either from the learned long-term-memory items or from the working-memory array; or neutral retrocues provided no information. The findings showed that internal attention conferred behavioral benefits for reporting contents of either origin. However, they also uncovered an interesting dissociation. Gaze measurements replicated the typical spatial biases when orienting attention to attributes of items encoded into working memory but, surprisingly, there was no evidence for spatial gaze biases when orienting attention to attributes of items from long-term memory.

In the current study, we sought to investigate further the mechanisms engaged by internal attention to similar contents with different origins by using high-temporal-resolution scalp electroencephalography (EEG) in the experimental design by Gong et al. (2025). Using EEG, we aimed to test the extent of similarity between the neural dynamics for selecting and prioritizing contents encoded into working memory or retrieved from long-term memory. One possibility is that similar neural mechanisms unfold, but that the spatial markers do not impact the oculomotor readouts in the case of long-term-memory contents. Another possibility is that the neural mechanisms differ significantly. Two different scenarios are compatible with this latter possibility. (a) Visual contents stored in long-term memory may acquire a different representational format and no longer require tethering to the spatial information available during learning. During selection and retrieval, the traces are reinstated into working memory, but are non-spatial in nature. (b) Alternatively, internal attention for contents in long-term memory may operate through distinct processes than those operating within recently encoded working memory, involving direct targeting of relevant traces in a content addressable way. The design is well suited for comparing the internal attention for the same type of memory (color and shape features of visual objects) under the same demand characteristics (reproducing object shapes with precision reports), thus equating motivational and other state variables within a common task.

## Methods

### Participants

The experimental procedures were reviewed and approved by the Central University Research Ethics Committee of the University of Oxford. A total of 30 human volunteers (22 females, 8 males; mean age = 24.67, *SD* = 6.22; 28 right-handed, 2 left-handed) with reported normal or corrected-to-normal vision were recruited. The sample size for the experiment was set before data collection, to match our prior study, which used a similar approach (Gong et al., 2025). Each participant provided written consent before participation and was reimbursed £20 per hour.

### Stimuli, task, and procedure

The main experimental task required participants to reproduce the shape of one of four colored shapes associated with the four quadrant locations. Two quadrant locations were associated with colored shapes learned during a previous learning session. For convenience, we refer to these as the long-term-memory (LTM) items. The two other quadrants were used to present items that were encoded afresh into working memory. We refer to these as working-memory (WM) items. Colored retrocues oriented attention to LTM or WM items or provided no information. The LTM and WM labels are intended to be neutral with respect to the state of these representations at the time of the retrocues. Contents retrieved from LTM could be stored as latent states or be reinstated into an active WM buffer (e.g., Cowan, 1999). Additionally, WM contents could include both active and silent representational states (e.g., Panichello et al., 2024; Stokes, 2015).

Participants sat in front of a 27ʺ monitor (1920 × 1080 pixels, 100 Hz) and rested their chin on a chinrest placed 95 cm away from the monitor. The experiment was programmed in MATLAB (MathWorks, Natick, MA, USA) using the Psychophysics Toolbox (Brainard, 1997). Stimuli appeared overlaid on a white background. Throughout the experiment, four squares (2.5° in diameter) were always presented as placeholders at the four quadrants, at 5° horizontally and vertically from the central fixation to the center of each square. Five equiluminant colors (burnt orange [205.5, 98.5, 29.5], olive green [111.5, 138, 61.5], teal blue [49, 140.5, 157], soft purple [170.5, 104, 185.5], and deep pink [245.5, 37, 112.5]) were drawn from a circle in CIELAB color space (center at *L\** = 54, *a\** = 18, *b\** = -8, radius = 59). For each participant, one of these colors was randomly assigned as the color of the neutral retrocue, and the remaining four were randomly paired with four shapes equidistantly sampled from the validated circular shape (VCS) space (Li et al., 2020). Out of the four pairs of colored shapes, two were randomly assigned as WM items, and the remaining two were used as LTM items. Each LTM item was consistently presented at the same diagonal location throughout the learning and testing sessions (e.g., always top-left and bottom-right for a given participant, counterbalanced across participants). WM items appeared within the complementary pair of diagonal squares. The experiment included a learning session and a testing session, separated by a 5-minute break.

During the learning session, participants were trained to memorize the colors and shapes of LTM items. Each learning trial began with a fixation display lasting randomly between 800 and 1000 ms, after which the LTM items were presented for 150 ms. Following a delay of 850 ms, one of the LTM locations was highlighted, and a color wheel or a shape wheel appeared at the center, indicating the feature to report in this trial. Both the color and shape wheels were presented in a random orientation each time. The shape wheel consisted of 360 shapes from the VCS space. To avoid clustering, eight shapes sampled from equidistant positions on the wheel were displayed along the cardinal axes (i.e., every 45 degrees). These eight shapes served as visual anchors, which were also randomly chosen every time based on the orientation of the shape wheel. By moving the dial, participants could inspect all intermediate shapes across the full 360-shape continuum in the VCS space. The color wheel likewise allowed continuous inspection across a full 360° color space. We adopted a continuous-report paradigm because it enables fine-grained measurement of reproduction precision and therefore beneficial for detecting subtle cue benefits on accuracy. Participants responded using a computer mouse that controlled the dial on the wheel. Participants had unlimited time to retrieve the content from memory and to decide what to reproduce. However, once they started moving the dial, they had only 2500 ms to complete their reproduction. This was intended to encourage participants to retrieve the exact color or shape before moving the dial. The position of the dial when participants clicked the left mouse button or when the time limit was reached was taken as the response. Immediately after their response, feedback was presented for 500 ms in the form of an integer ranging from 0 to 100, with 100 indicating a perfect reproduction of the probed color/shape and 0 indicating the exact opposite on the color/shape wheel. Each color and shape were probed on 20 trials, resulting in a total of 80 learning trials presented in random order.

During the testing session, participants were presented with the WM items on every trial and performed an LTM/WM shape reproduction task. Each testing trial began with a fixation display (randomized between 800-1000 ms), after which the WM items were presented for 150 ms. The color-shape bindings of WM items were randomly swapped from trial to trial. Following a delay of 850 ms, a color retrocue was presented for 200 ms. On one-third of the trials, the retrocue was neutral, and participants were probed with one of the four LTM/WM colors following another delay of 800 ms. On the remaining two-thirds of the trials, the retrocue was informative, matching one of the LTM/WM colors with equal probability, and participants reproduced the shape of the cued item following the second delay. The shape wheel was identical to that used during the learning session and was randomly rotated across trials. The testing session consisted of 480 trials divided into 10 blocks (each including 48 trials). To become familiarized with the task, participants performed an additional 20 practice trials before testing.

### Behavioral analysis

Reproduction errors were defined as the absolute difference (in degrees) between the reported color/shape and the actual color/shape of the target item. Response time was defined as the time from probe onset to report initiation (i.e., start moving the dial).

All learning trials were kept for analysis. We examined the learning performance by sorting color and shape learning trials into 4 bins (each containing 10 trials) and submitting the reproduction errors to one-way repeated-measures ANOVAs with linear contrast weights ([−3, −1, 1, 3]) across 4 bins.

In the testing session, trials with a response time beyond 3 SD of the individual mean were excluded. Overall, 2.0% ± 1.3% (M ± SD) trials were excluded after this step. To compare behavioral performance between testing conditions, we submitted reproduction errors and response times to repeated-measures ANOVAs and reported partial *η*^2^ as a measure of effect size. Where relevant, the within-subject standard error of the mean was calculated from normalized data (Morey, 2008). For visualization, the probability densities of response times were fitted using lognormal distributions, as response times typically exhibit positive skewness characteristic of this distribution (Thissen, 1983; van der Linden, 2006). Individual lognormal distributions were fitted to each participant’s RT data for each condition using maximum likelihood estimation. We then fitted lognormal distributions to pooled RT data across all participants within each condition to characterize the overall RT distribution shape. Reproduction error densities were fitted using kernel density estimation (KDE) to avoid assumptions about the underlying distribution shape, as error distributions may exhibit complex patterns including multi-modality or non-standard skewness. Individual KDE curves were fitted to each participant’s reproduction error data using Gaussian kernels with adaptive bandwidth selection, and then we fitted KDE curves to pooled data across all participants within each condition to characterize overall error distribution shapes.

### Eye tracking acquisition and analysis

Eye movements were recorded with an EyeLink 1000 Desktop Mount (SR Research, Ottawa, ON, Canada) positioned ∼15 cm in front of the monitor. Before recording, the eye tracker was calibrated through the built-in calibration and validation protocols from the EyeLink software. Horizontal and vertical gaze positions were continuously recorded for both eyes at a sampling rate of 1000 Hz. For some participants, only one eye was tracked due to a lack of good-quality binocular tracking (n = 3). Data were first converted from edf to asc format and subsequently read into RStudio. For binocularly tracked participants, data from the left and right eyes were averaged to obtain a single horizontal and a single vertical gaze-position channel. Blinks were marked by detecting NaN clusters in the eye-tracking data. The data during 100 ms before and after detected NaN clusters were rewritten as NaN to eliminate residual blink artefacts. Finally, data were epoched from −200 to 1000 ms around cue onset and baseline corrected by subtracting the average gaze position during the 200-ms window preceding cue onset. Gaze time courses were then smoothed using a 20-ms average moving window. Following previously described methods (Draschkow et al., 2022; Gong et al., 2025; van Ede, Chekroud, & Nobre, 2019), we focused our analyses on the horizontal channel and only included trials in which gaze position remained within ±50% from fixation (with 100% denoting the centers of the original item locations at a ±5° visual angle) throughout the trial. On average, 3.5% ± 4.4% (M ± SD) of the trials were excluded from each participant.

Task modulations of gaze position were first identified by comparing trial-averaged gaze position time courses between conditions in which the retrocued item occupied the left side (top/bottom left location) or the right side (top/bottom right location) during encoding, separately for trials with WM retrocues and trials with LTM retrocues. To increase sensitivity, we constructed a measure of towardness separately for trials with WM and LTM retrocues, which expressed the gaze bias toward the side of the retrocued item in a single value. Trial-averaged gaze-position time courses were also obtained for trials with neutral retrocues.

For visualization purposes, we constructed a heatmap of gaze positions between 400 and 800 ms following retrocue onset, averaged across all retrocue conditions. For this analysis, we did not remove trials with gaze values exceeding half the distance to memory item locations. Two-dimensional kernel density estimations were obtained separately for trials where retrocued items occupied the left/right side. Lastly, we subtracted left-cued gaze density maps from right-cued gaze density maps to illustrate the magnitude of the observed gaze bias.

### EEG acquisition and preprocessing

EEG data were acquired using a SynAmps amplifier and Curry 8 acquisition software (Compumedics NeuroScan, Charlotte, NC, USA). Sixty-one electrodes were distributed across the scalp following the international 10–20 system. An electrode placed behind the left mastoid was used as the active reference during acquisition. The ground was placed at the middle of the forehead. Two bipolar electrode pairs recorded electrooculography (EOG); one above and below the left eye (vertical EOG) and another lateral to each eye (horizontal EOG). Data were digitized at 1,000 Hz and low-pass filtered at 500 Hz during acquisition.

Offline, EEG data were analyzed using MNE-Python (Gramfort et al., 2013). First, data were high-pass (0.1 Hz) and low-pass (60 Hz) filtered with a Finite Impulse Response (FIR) filter and downsampled to 250 Hz. Then, data were re-referenced to the common average. Bad electrodes were manually identified and interpolated. On average, 2.03 ± 1.85 (M ± SD) bad electrodes were identified and interpolated per participant. Data were then epoched from −200 to 1000 ms relative to the retrocue onset. An independent component analysis (ICA) and an automatic labeling algorithm (Pion-Tonachini et al., 2019) were used to remove artifacts associated with eye movements, muscle artifacts, heartbeats, and channel noises. ICA was fitted separately on 1-Hz high-passed data and then applied to the original 0.1-Hz high-passed data. We also ran the autoreject (local) algorithm (Jas et al., 2017) to interpolate or exclude noisy epochs. In total, 94.68 ± 5.46% of trials were retained for subsequent analyses.

### Time-frequency analysis

Time-frequency representations of the EEG data were computed using Morlet wavelet convolution. Morlet wavelets were constructed for frequencies ranging from 3 to 60 Hz in 1-Hz steps. The number of cycles for each frequency was set to half the frequency, balancing temporal and spectral resolution. The wavelet transform was applied to each epoch using Fast Fourier Transform (FFT) for computational efficiency. The resulting power estimates were then baseline-normalized using a percent change relative to the 200-ms pre-stimulus interval. To investigate the lateralization of neural activity relative to the memorized location of the cued item, we focused our analyses on canonical left and right visual electrodes (PO7 for left visual, PO8 for right visual). Time-frequency representations were contrasted between trials in which the cued WM/LTM item was contralateral versus ipsilateral to the electrode of interest. We expressed this contrast as the difference of percentage change in power (contra minus ipsi) and averaged it across the left and right visual electrodes. Topographical maps of alpha lateralization were obtained by applying the same procedure to all symmetrical electrode pairs (left hemisphere: ipsi minus contra, right hemisphere: contra minus ipsi). To derive time courses of alpha lateralization, we averaged the contralateral versus ipsilateral response within the predefined alpha band (8–12 Hz). We also ran the same procedures on EOG-regressed EEG data to make sure our time-frequency analysis was not contaminated by eye movements.

### Multivariate pattern analysis

To examine whether neural activity patterns differed between conditions and how the differences evolved over time, we implemented a temporal decoding analysis to perform classification at each time point independently. This method trains separate classifiers for each time point, enabling the assessment of when neural activity patterns differ between conditions. First, features (i.e., evoked responses across sensors at a certain time point) were standardized using z-score normalization. Second, principal component analysis (PCA) was applied with 99% variance retention to reduce dimensionality while preserving the majority of the signal variance. Finally, linear discriminant analysis (LDA) was used as the classification algorithm to find the optimal linear combination of features that maximally separated the experimental conditions. Three-fold cross-validation was implemented within each time point to ensure robust generalization. This approach randomly splits the data into three folds, training the classifier on two folds and testing on the remaining fold, repeating this process three times with different fold assignments. Classification performance was assessed using the Area Under the Curve of the Receiver Operating Characteristic Curve (ROC-AUC), which is considered a robust and criterion-free measure of classification (Hand & Till, 2001; Thölke et al., 2023). ROC-AUC values range from 0.5 (chance performance) to 1.0 (perfect discrimination). Lastly, the trial-averaged decoding time courses were smoothed using a Gaussian filter with a 20-ms SD.

In addition to temporal decoding, we implemented temporal generalization analysis (Grootswagers et al., 2017; King & Dehaene, 2014) to assess the temporal stability and persistence of neural representations across time. This approach tests whether neural patterns that are discriminable at one time point remain discriminable at other time points, providing insight into the temporal dynamics of information processing. Specifically, by training classifiers at each time point and testing them at all time points, this approach creates temporal generalization matrices in which each element (i, j) represents the classification performance of a classifier trained at time point i and tested at time point j.

To test which brain areas contributed most to the classifier likelihoods observed in our multivariate methods, we ran a sensor-space searchlight decoding analysis. For this analysis, we ran the same analysis as outlined above across small clusters of neighboring EEG sensors, resulting in decoding scores for each sensor and time point. These decoding scores were averaged into time clusters of 200 ms each, resulting in 5 topographical maps between 0 and 1000 ms. These topographical plots show whether decoding results were primarily driven by specific sensor clusters. To evaluate the statistical significance of sensor clusters with decoding scores above chance level, we performed cluster-based permutation tests, which took account of the adjacency matrix for the 61 EEG sensors.

To test for potential eye-movement contributions towards decoding findings (Mostert et al., 2018; Quax et al., 2019), besides conducting the analysis on EEG data, we ran the same procedures on another two datasets: EOG data and EOG-regressed EEG data. Comparing the decoding scores from these two sets of analyses with the original one then allows us to assess the impact of eye movements on the EEG decoding results.

To control for potential confounds related to low-level stimulus features, we implemented a cross-decoding analysis to test whether a classifier could generalize across different color cues when distinguishing WM from LTM retrocue trials. For this analysis, the two WM-associated colors and two LTM-associated colors were grouped into two pairs. We trained an LDA classifier on trials corresponding to one pair of colors (WM color 1 vs. LTM color 1) and tested it on the held-out trials from the second pair (WM color 2 vs. LTM color 2). All other aspects of the analysis pipeline remained identical to the primary decoding analyses.

### Relationship between decoding and behavior

To investigate the relationship between the observed decoding of WM vs. LTM retrocue and the behavioral performance, we employed the following approach. Firstly, we computed timepoint-by-timepoint decoding scores of each trial and then averaged across time to derive a single score for each trial. Subsequently, we performed a median split based on these trial scores, creating two groups of trials: one with low decoding scores and the other with high decoding scores. We then compared the response times and reproduction errors of WM and LTM retrocue trials based on the median split.

### Statistical evaluation of time series

Statistical evaluation of time series was conducted using a cluster-based permutation approach (Maris & Oostenveld, 2007), which evaluates temporal clusters observed in the original data against a permutation distribution derived from the largest temporal cluster found after each permutation of the condition labels. Time series examined included oculomotor towardness time courses, the contralateral-versus-ipsilateral time-frequency spectra, the alpha attenuation time courses, and the decoding score time courses.

To investigate the temporal relationship between orienting to WM and LTM representations, we performed a cross-correlation analysis on the time-resolved decoding scores. Decoding score time series were first z-score normalized to ensure the cross-correlation reflected temporal relationships independent of overall amplitude differences. The cross-correlation z at lag k is computed as below:

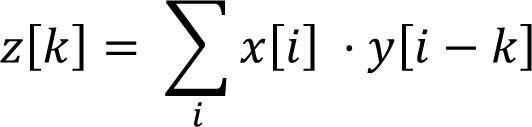

where x[i] represents the value of the LTM decoding score at index i, and y[i−k] represents the value of the WM decoding score at index i−k. Cross-correlations are computed across all possible temporal lags and were normalized by the length of the time series to ensure the values fall within the range [−1, 1]. This analysis allows us to assess the consistency of temporal relationships between orienting to WM and LTM neural representations. If the time course of WM decoding score precedes the time course of LTM decoding score, the maximum cross-correlation should occur at a positive temporal lag. For each participant, we determined the lag corresponding to the maximum cross-correlation coefficient and then compared them against zero.

## Results

The first experimental ingredient was for participants to learn the featural attributes of objects that would later serve as LTM items in the subsequent phase of testing. Participants first memorized the colors and shapes of two items presented at two diagonal locations during a training session (Fig. 1b). The learning session resulted in good color and shape reproduction over 4 trial bins (Fig. 1c). Color reproduction yielded high accuracy through the learning session. Errors remained at a relatively low level and decreased numerically across bins (first bin: 18.803 ± 2.285, last bin: 17.637 ± 1.633), although the improvement was not significant (*F*(1,29) = 0.341, *p* = 0.560). Shape reproduction improved significantly across bins (*F*(1,29) = 19.588, *p* < 0.001, partial η^2^ = 0.142).

**Figure 1.**
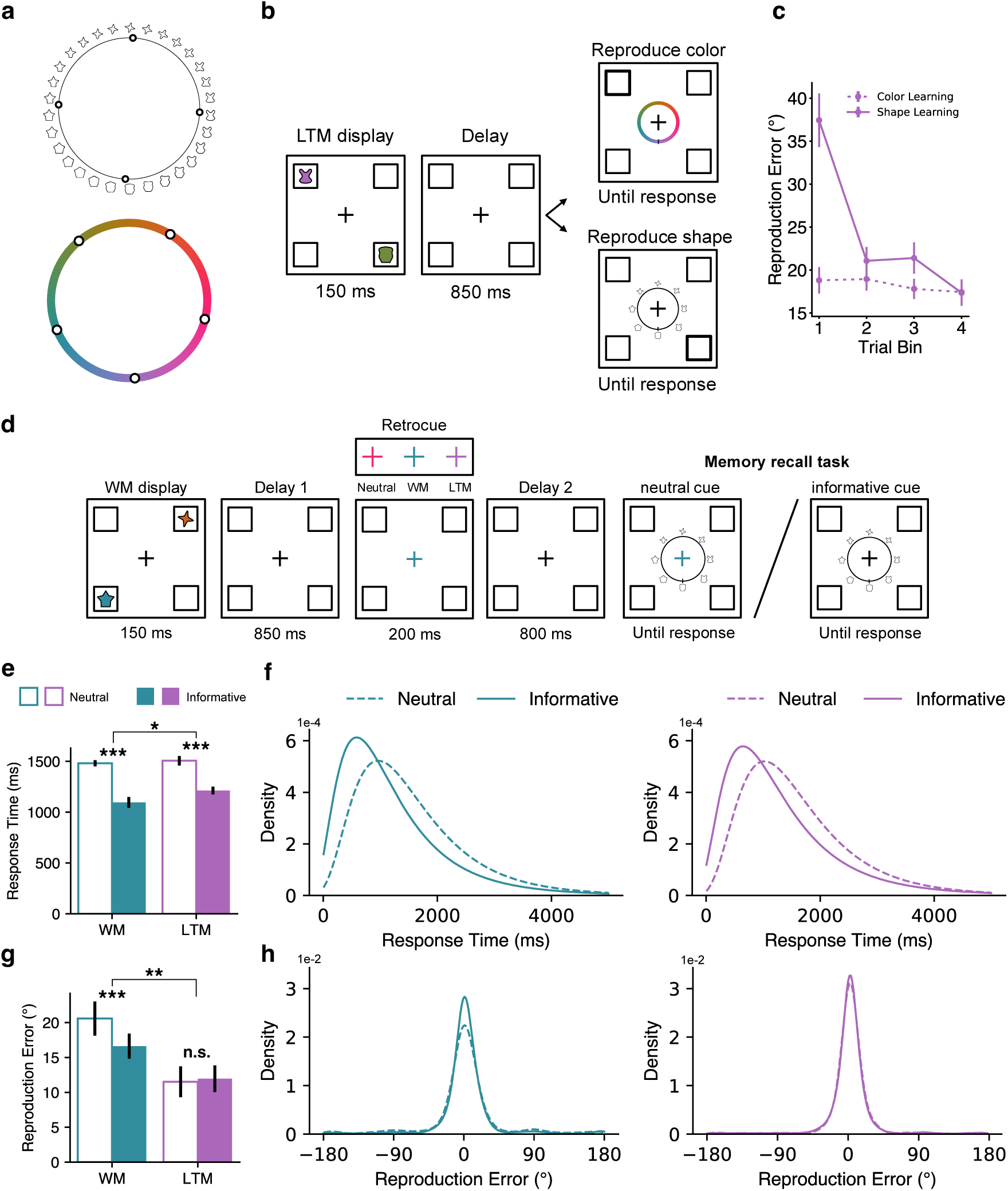
Task design and behavioral results. (a) 4 shapes sampled from the validated shape space and 5 colors sampled from a CIELAB color space were used as stimuli, forming 4 colored shapes (randomly assigned to WM and LTM) and one extra color as the neutral cue. (b) During the learning session, participants were trained to memorize the precise features of two LTM items by reproducing the color or shape of either item on each trial. (c) Participants exhibited improvements in color and shape reproduction (sorted into 4 trial bins). (d) During the testing session, participants first encoded two WM items. After a short delay, a color retrocue was presented at the fixation cross which can be neutral or informative (matching the color of one WM/LTM item). After a second delay, participants either reproduced the shape of the item matching a randomly chosen color (presented at the fixation cross) if the retrocue was neutral or reproduced the shape of the color-matching item if the retrocue was informative. On informative-cue trials, participants knew what to report after the retrocue; on neutral-cue trials, the item to report was revealed only at the response phase. (e) Response times for WM and LTM reproduction as a function of neutral and informative retrocues. (f) Probability densities of response times fitted to lognormal distributions (left panel: WM, right panel: LTM). (g) Reproduction errors of WM and LTM items as a function of neutral and informative retrocues. (h) Probability densities of reproduction errors were fitted using kernel density estimation. Error bars in (c), (e), and (g) represent ±1 SEM. *, **, ***, and n.s. indicate p < 0.05, p < 0.01, p < 0.001, and not significant, respectively.

### Orienting attention to WM items elicited stronger behavioral benefits than to LTM items

Consistent with results from the previous study (Gong et al., 2025), informative retrocues exerted significant main effects on both response times (*F*(1,29) = 302.973, *p* < 0.001, partial η^2^ = 0.913; Fig. 1e) and reproduction errors (*F*(1,29) = 10.502, *p* = 0.003, partial η^2^ = 0.266; Fig. 1g). Interaction effects showed that benefits were stronger for WM items, with a bigger reduction on both response times (*F*(1,29) = 5.410, *p* = 0.027, partial η^2^ = 0.157) and reproduction errors (*F*(1,29) = 5.659, *p* = 0.024, partial η^2^ = 0.163) following informative retrocues. LTM retrocues benefitted response times (*t*(29) = 7.362, *p* < .001, Cohen’s *d* = 0.218) but did not reduce reproduction errors significantly (*t*(29) = 0.598, *p* = .554). WM retrocues benefitted both response times (*t*(29) = 11.817, *p* < .001, Cohen’s *d* = 0.336) and reproduction errors (*t*(29) = 3.960, *p* < .001, Cohen’s *d* = 0.395). Overall, reproduction of LTM items was slower (*F*(1,29) = 7.854, *p* = 0.009, partial η^2^ = 0.213) but more accurate (*F*(1,29) = 7.931, *p* = 0.009, partial η^2^ = 0.215) compared to WM items. To confirm the effects above, we plotted probability densities of response times (Fig. 1f) and reproduction errors (Fig. 1h). These demonstrated the effects of informative retrocues and their differential pattern of benefits on WM and LTM reproduction.

### Gaze was biased toward memorized locations for WM but not LTM items

Analysis of eye-tracking data replicated the oculomotor dissociation discovered in the previous study (Gong et al., 2025). When WM items were retrocued, gaze was biased toward the location (left vs. right) where the cued item was encoded. In contrast, when LTM items were retrocued, gaze stayed near baseline (Fig. 2a). Towardness time courses, computed by combining right- and left-cued gaze time courses to compute systematic gaze shifts toward vs. away from the cued item, confirmed that retrocues biased gaze significantly in the direction of attentional orienting to WM items (cluster *p* = 0.001), while no statistically significant clusters were identified when orienting to LTM items (Fig. 2b). The difference in gaze biases between WM and LTM conditions was significant (cluster *p* = 0.001). The density map computed by subtracting left-cued WM trials from right-cued WM trials demonstrated that this bias was primarily composed of small gaze shifts toward the location of the cued WM item (Fig. 2c), in line with previous findings (Draschkow et al., 2022; Gong et al., 2025; van Ede, Chekroud & Nobre, 2019).

**Figure 2.**
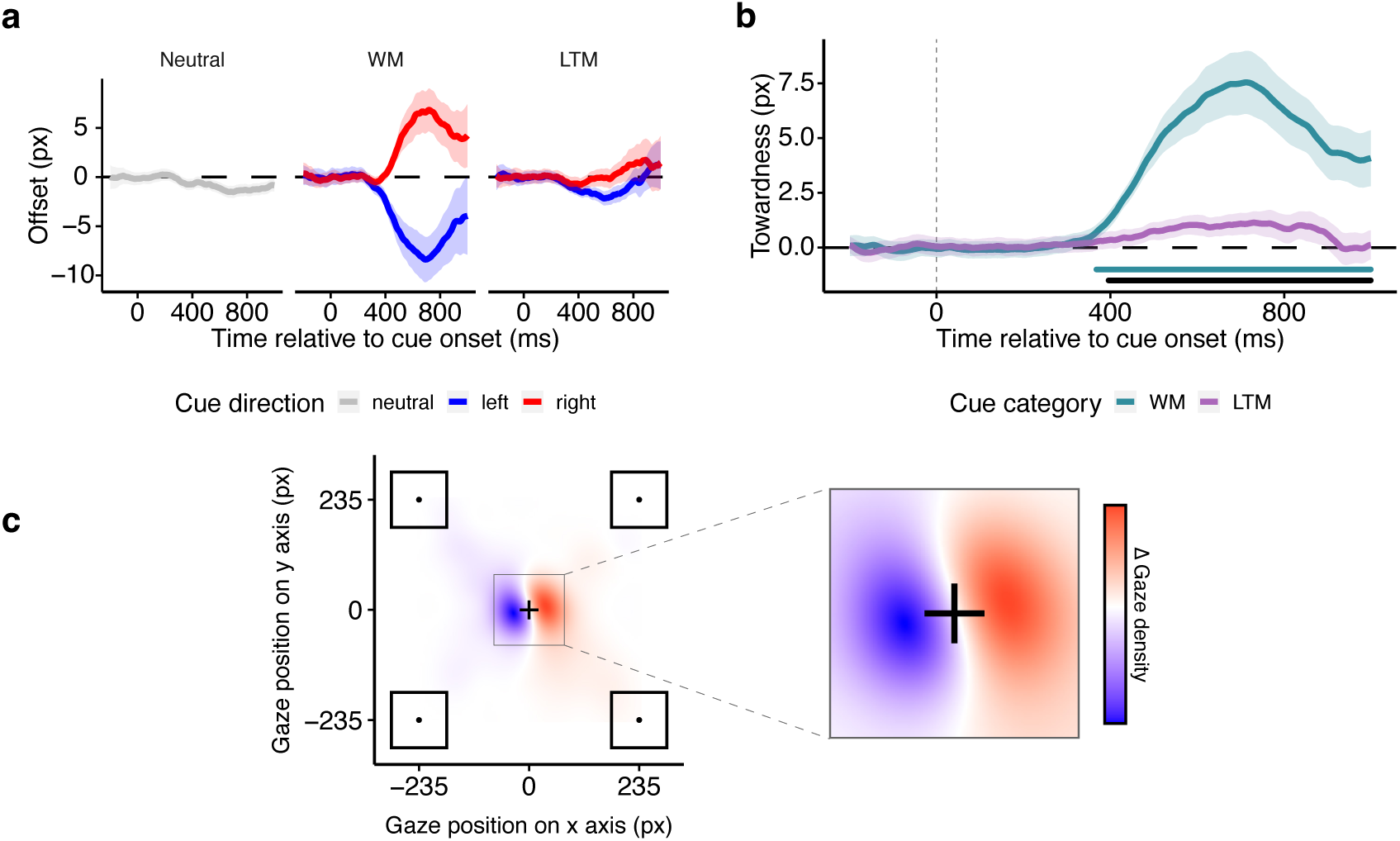
Informative retrocue biases gaze toward cued WM items but not LTM items. (a) Time courses of horizontal gaze position as a function of cue domain and cue direction. (b) Time courses of average horizontal gaze biases expressed as “towardness” for WM and LTM retrocue trials. Horizontal lines below the time courses indicate significant temporal clusters (teal blue: WM compared to 0; black: WM vs. LTM difference). (c) Difference in gaze density following WM retrocues (right vs. left), obtained by subtracting the density map of left-cued trials from the density map of right-cued trials over the time window of 400-800 ms after cue onset. Shading areas in (a) and (b) represent ±1 SEM.

### Alpha attenuation was observed during WM selection, but not LTM selection

The contralateral suppression of posterior 8- to 12-Hz alpha activity is a canonical marker of spatial attention shifts both in perception (Doesburg et al., 2016; Gould et al., 2011; Jensen & Mazaheri, 2010; Rihs et al., 2007; Worden et al., 2000) and working memory (Myers et al., 2015; Poch et al., 2014; van Ede, 2018; Wallis et al., 2015; Woodman et al., 2022). In our experimental design, WM and LTM items were memorized at two pairs of lateral but non-overlapping locations, allowing us to compare alpha attenuation (contralateral vs. ipsilateral) during the selection of WM and LTM contents. When orienting to WM items, alpha attenuation emerged during the second half of delay period (cluster *p* = 0.033) and occurred at posterior electrodes (Fig. 3a). However, no contralateral alpha suppression was found when orienting to LTM items (Fig. 3b). We also directly compared the time courses of contralateral vs. ipsilateral alpha power (averaged over 8 to 12 Hz; see Fig. S2 for the evolution of corresponding topographies averaged in 200-ms steps). The comparison confirmed that there was significant lateralized alpha attenuation following retrocues to WM items (cluster *p* = 0.009), which was significantly more pronounced than in the LTM condition (cluster *p* = 0.034). Although it is clear from the topography that alpha attenuation mainly occurred in the posterior region, we ran the same analysis on EOG-regressed EEG data and confirmed that this effect was not just explained by eye movements (Fig. S1).

**Figure 3.**
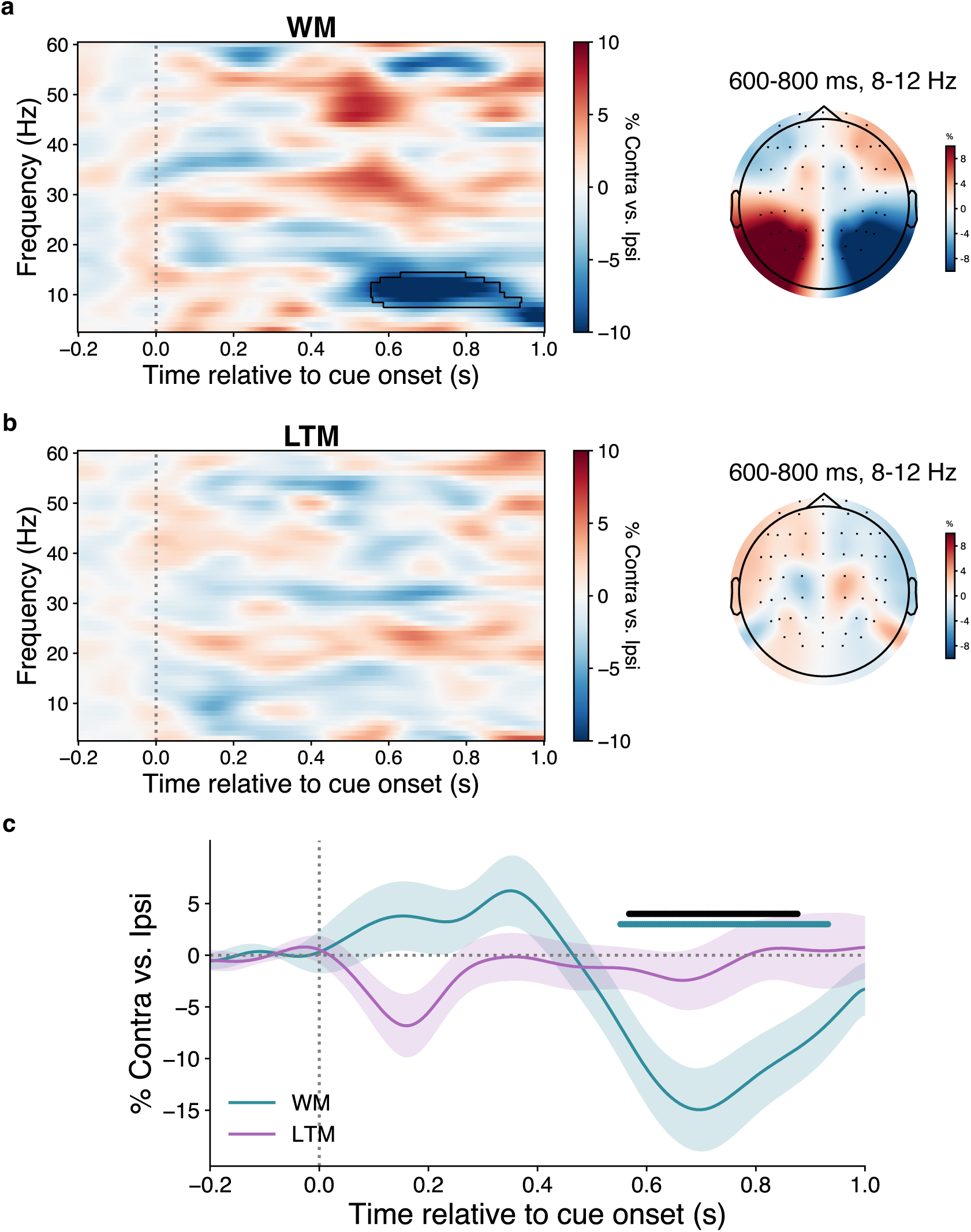
Lateralized neural activity following retrocues to WM and LTM items. (a) Left: normalized time-frequency difference calculated as percent change in power contralateral vs. ipsilateral to the memorized location of the cued WM item in visual electrodes PO7/PO8. Right: topography representing time-frequency difference across all sensor pairs averaged over 600-800 ms and 8-12 Hz. The two halves of the topography are symmetrical, representing ipsilateral minus contralateral (left hemisphere) and contralateral minus ipsilateral (right hemisphere), respectively. Black outlines in the time-frequency plot indicate significant clusters compared to 0. (b) Same as (a) but for retrocues to LTM items. (c) Time courses of alpha lateralization in the alpha band (8-12 Hz) represented by contralateral minus ipsilateral power difference. Horizontal lines represent significant clusters (teal blue: WM compared to 0; black: WM vs. LTM difference). Shading areas represent ±1 SEM.

### Orienting attention to WM and LTM evoked distinct neural activity patterns

To investigate whether different neural processes underlie attentional orienting to WM and LTM contents, we applied time-resolved multivariate classification using linear discriminant analysis (LDA). The broadband EEG signal was used to decode the cognitive states associated with selection of WM and LTM items. This allowed us to track the evolution of neural discriminability between focusing on WM and LTM items at high temporal resolution. Fig. 4a (upper panel) shows robust decoding of informative-retrocue trials relative to neutral-retrocue trials (WM items: cluster *p* = 0.001; LTM items: cluster *p* = 0.003 (see Fig. S3 for additional generalization-across-time analyses). Moreover, selecting contents from WM items led to more robust decoding than selecting contents from LTM items (cluster *p* = 0.003). To identify the electrodes that contributed more strongly to the decoding scores, we conducted a searchlight analysis with subsets of EEG sensors used iteratively in decoding (as in Kriegeskorte et al., 2006; van Ede, Chekroud, Stokes, et al., 2019). Cluster-based permutation tests accounting for electrode adjacencies showed that orienting attention to WM items engaged a more widespread topography compared to LTM items, with more sensors showing decoding scores above chance level (Fig. 4a lower panel). Because overall decoding strength can influence how many electrodes surpass a fixed threshold during permutation tests, we also trained a classifier to directly discriminate WM trials from LTM trials to help interpret contrasts between the two conditions (Fig. S4).

**Figure 4.**
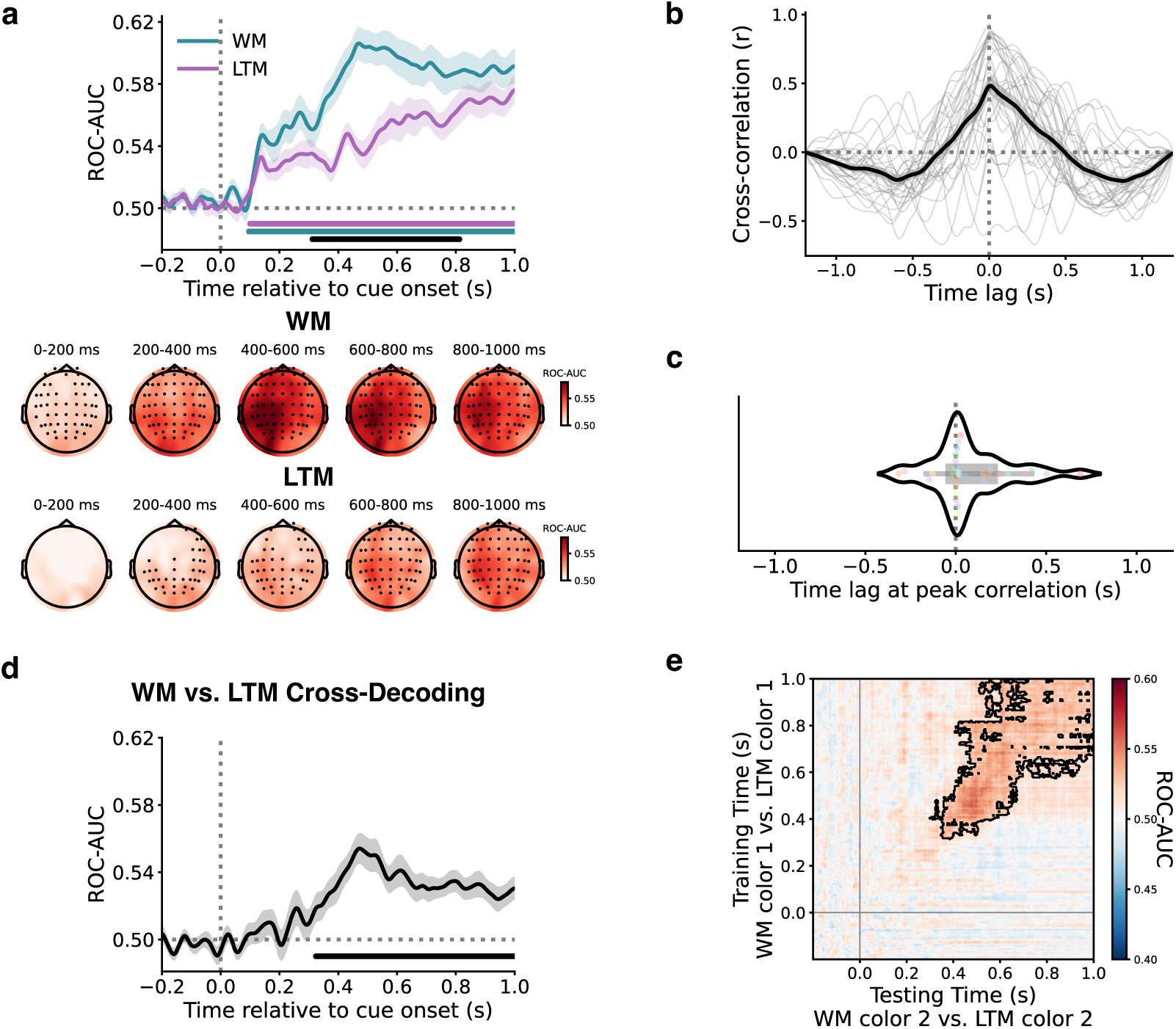
Time-resolved EEG decoding of attentional orienting in WM and LTM representations. (a) Upper panel: decoding score time courses of WM (vs. neutral) and LTM (vs. neutral) conditions, time-locked to cue onset. Lower panel: decoding topographies computed using a searchlight approach, which iteratively trained the classifier on each electrode and its immediate neighbors to get decoding scores for each electrode and time point. Topographies were averaged in 200-ms steps. (b) Cross-correlation coefficients between the time courses of WM and LTM decoding scores in (a). Gray lines represent individual participants. (c) Time lags at which cross-correlation coefficients reached their maximum. Colored dots represent individual participants. (d) Decoding score time course of the classifier trained to distinguish trials with one pair of item colors (WM color 1 vs. LTM color 1) and tested on trials with the other pair of colors (WM color 2 vs. LTM color 2). (e) Temporal generalization matrix of the classifier performance in discriminating WM and LTM conditions independent of specific color cues. The diagonal corresponds to the time course in (d). Black outlines in the matrix indicate significant clusters of temporal generalization above chance level. Shading areas in (a), (b), and (d) represent ±1 SEM. Horizontal lines in (a) and (d) indicate significant temporal clusters (teal blue: WM condition compared to 0; soft purple: LTM condition compared to 0; black: difference between WM and LTM conditions. Black dots on topographies in (a) and (d) indicate significant electrode clusters with decoding scores above chance level.

To compare the temporal dynamics of modulations when focusing attention on LTM versus WM items, we cross-correlated the respective decoding time courses. The analysis revealed that the maximal correlation consistently occurred at a near-zero time lag (Fig. 4b). The finding was substantiated by a direct comparison of the peak correlation time lags against zero (Fig. 4c), which did not reveal a significant difference (*t*(29) = 1.928, *p* = 0.064, Cohen’s *d* = 0.352). This temporal alignment suggests that the neural processes initiated by retrocues unfold over a similar timescale for both WM and LTM items.

To ensure that the classifier was discriminating between the cognitive states of orienting to WM vs. LTM items independently of any physical properties of the color cues, we ran a cross-decoding analysis. We trained a classifier to distinguish between orienting to WM and LTM items using one pair of item colors (WM color 1 vs. LTM color 1) and tested its performance on trials with the other pair of colors (WM color 2 vs. LTM color 2). This analysis revealed robust cross-decoding (cluster *p* = 0.001), with classification performance rising significantly above chance around 300 ms after cue onset (Fig. 4d). The cross-decoding analysis establishes that different patterns of neural activity are associated with selecting WM vs. LTM items, independently of the specific sensory qualities of the cues. The later onset of pattern differences relative to original decoding analyses suggests that the earliest pattern separation between WM and LTM conditions in the original analyses may have reflected sensory processing of the retrocue stimulus (Fig S4). The temporal generalization matrix of the cross-decoding analysis (Fig. 4e) reveals how the stability of this representation evolves. Over time, the pattern changes from more dynamic to more stable.

Eye movements may contaminate EEG recordings and lead to decodable non-brain signals (Mostert et al., 2018; Quax et al., 2019). Therefore, the analyses were repeated on EOG data and on EOG-regressed EEG data to test whether our decoding results were driven by eye movements. All decoding results remained robust after EOG activity was regressed out (Fig. S5a-c), and eye movements contributed little to decoding scores.

In an exploratory analysis, we tested whether patterns of brain activity discriminating neutral retrocues from informative retrocues to WM items or LTM items impacted behavioral performance (Fig. S6). Single-trial decoding scores averaged across all time points were derived for each participant and categorized into low vs. high decoding performance. The results showed that high decoding performance predicted both shorter response times (*p* = 0.025) and lower reproduction errors (*p* = 0.028) when orienting to WM items. Response times (*p* = 0.568) and reproduction errors (*p* = 0.431) did not differ for LTM items with low vs. high decoding performance.

## Discussion

The present study provides converging behavioral, oculomotor, and neural evidence that internal attention processes differ when operating on contents from WM vs. LTM to retrieve the same information (visual shape) within the same task. The two types of attention processes were engaged flexibly, on a trial-by-trial basis, depending on which item was relevant and its domain of origin. The findings add interesting new insights for understanding internal attention.

Orienting internal attention to reproduce the shape of an item yielded performance benefitted whether the item was just encoded into WM from the sensory stream or had been learned in a previous session and stored in LTM. The pattern of behavioral facilitation differed based on the memory domain of origin and replicated previous observations (Gong et al. 2025). Accuracy benefits were exclusive, and response-time benefits were stronger, for WM than LTM items.

Gaze and neural recordings showed that these behavioral benefits involved different mechanisms. Eye-tracking results replicated the oculomotor dissociation reported in the previous study (Gong et al., 2025): orienting attention to a WM item induced a reliable gaze bias toward the cued item’s location, replicating previous findings (Draschkow et al., 2022; van Ede, Chekroud, & Nobre, 2019). In contrast, the signature was absent when orienting to an LTM item. Thus, oculomotor signals indicated stronger reliance on spatial scaffolding for orienting internal attention to WM items (see Liu, Nobre, & van Ede, 2022) than LTM items in this task.

The EEG results extended the observation of greater reliance on spatial mechanisms when orienting internal attention to WM items than LTM items. The well-established neural marker of spatial attention, the contralateral suppression of posterior alpha-band activity, was clearly present when retrocues oriented attention to WM items but was absent for LTM items. As in previous WM studies, the spatial modulatory mechanism occurred when orienting to WM items even though item location was not necessary for the shape reproduction task. The findings support many observations of the importance of spatial coding to scaffold attention and access to recently encoded WM contents, even when there are no specific spatial demands in the task (Curtis et al., 2022; Foster et al., 2017; Pertzov & Husain, 2014; Schneegans & Bays, 2017; van Ede et al., 2017).

The different extent of reliance on spatial processes when orienting internal attention to WM and LTM items is revealing in at least two ways. At a more general level, it indicates that there are different types of mechanisms for visual internal attention, one that invokes spatial processes even when spatial information is not required for task performance, and one that can operate without robustly engaging canonical spatial markers. We are not aware of other studies reporting selective modulation of internal visual representations from competing items in an array in the absence of spatial mechanisms.

The finding also sheds light on the processes of orienting internal attention to items originating from LTM. They indicate that the lack of evidence for gaze biases when orienting to LTM items reflects differences in the neural processes engaged during internal attention orienting. Interestingly, there was no evidence for lateralized alpha modulation when orienting to LTM items despite the LTM feature associations being originally acquired by probing features by their locations, thus relying on strong spatial contexts. One caveat is that the absence of spatial EEG markers for LTM does not guarantee that no spatial mechanisms were deployed. Spatial mechanisms could have gone undetected due to methodological limitations. However, even accounting for this, the overall results show strong and significant differences in the type or extent of spatial mechanisms engaged when orienting internal attention to WM vs. LTM contents.

This interpretation is bolstered by the multivariate decoding analysis. The direct comparison of neural patterns when orienting to WM vs. LTM items revealed that the two conditions were distinguishable for a sustained period, over and above any effects of retrocue colors.

The distinct patterns of brain activity engaged by internal attention to WM and LTM items have alternative possible explanations. In one scenario, both items encoded from the sensory stream and those retrieved from LTM are placed and maintained in a WM state, but they have different properties depending on their origin. In this “representational-formats” account, internal attention operates upon different types of content within the same, WM domain. In the alternative scenario, items encoded from the sensory stream and those stored in LTM may exist in different memory domains, with internal attention operating distinctly upon the different “representational domains”. By either account, the processes involved in internal attention were shown to vary flexibly between more and less spatial mechanisms depending on the origin of the attended item.

Under the representational-formats account, consolidated LTM representations may have shifted from detailed perceptual codes toward more abstract formats emphasizing the relevant featural dimensions of the stimuli (Hitch et al., 1995; Huang et al., 2024; Lifanov et al., 2021). If LTM traces are represented in a less spatially dependent format, spatial orienting mechanisms may not need to be engaged to support selective retrieval. This interpretation is compatible with findings that repeated retrieval and consolidation can transform memory representations toward more schematic, feature-based formats (Lifanov et al., 2021).

Under the representational-domains account, the distinct pattern of modulation associated with internal attention for WM and LTM items could represent dissociable mechanisms for selecting and prioritizing internal contents originating from the sensory stream or LTM. For example, not finding evidence for alpha lateralization could point to the existence of mechanisms for influencing LTM contents that sidestep the posterior parietal-frontal spatial attention pathways involved in WM control, which are closely related to oculomotor functions (Nobre et al., 1994).

The present data alone cannot distinguish between these accounts. The cross-correlation analysis showed a zero-lag relationship between the onset of decoding in the WM and LTM conditions, indicating that orienting to LTM items is not simply a delayed version of orienting to WM items. This finding is informative but not decisive with respect to the two accounts. Under the distinct representational-domains account, the zero-lag would reflect parallel but separate selection processes operating in different memory domains, which happen to unfold on similar timescales. Under the representational-format account, similar selection processes would operate within the same domain but target different attributes, depending on differences in representational formats and their reliance on spatial codes.

What the zero-lag finding rules out is the simplest version of a serial gateway model, in which LTM contents must first be reinstated in WM at the time of the retrocue before selection can begin. Such a model would predict a temporal lag with brain activity related to WM selection appearing later on LTM selection trials. However, more nuanced versions of a WM-mediated account, in which transfer is rapid or concurrent with retrieval, remain compatible with the observed temporal alignment.

The central contribution of the present work is to demonstrate that the neural profile of internal orienting is not unitary: spatial attention markers are clearly and robustly different when attention is directed to WM versus LTM items in our task. Further experiments will be required to clarify whether the attributes of items recently encoded into WM from the sensory stream and retrieved from LTM are represented within separate domains or a common domain. Furthermore, if LTM items are reinstated into WM, they may exist in an equivalent state as WM items or have a characteristic state and display particular properties.

Another important avenue for future research is identifying the neural pathways that support internal attention to LTM items in the absence of typical spatial selection markers. The absence of posterior alpha suppression suggests a departure from the typical top-down perceptual modulation by the dorsal frontoparietal network (Nobre et al., 2004; Szczepanski et al., 2013; Wallis et al., 2015). Future studies combining EEG with fMRI could help localize the sources of alternative, non-spatial selection signals. Additionally, it will be interesting to explore the boundary conditions of this dissociation. How does the nature of the encoded information (e.g., spatial vs. non-spatial features), cues (e.g., directional vs. associative cues), and task demands (e.g., reproduction vs. discrimination) influence which attentional processes are engaged?

In conclusion, our findings provide convergent evidence that the markers of internal attention differ markedly when orienting to WM versus LTM contents. Attention to WM contents is strongly grounded in spatial processing mechanisms, characterized by contralateral alpha suppression and overt gaze biases. In contrast, orienting to LTM contents in our task proceeded without these markers of spatial attention. Whether this dissociation arises because consolidated LTM representations lack the spatially detailed format that engages spatial gating, or because selection of LTM items recruits fundamentally different selection mechanisms, remains an important open question. Resolving it will require future designs that can independently manipulate the representational formats and memory stores. What the present results establish is that the neural profile of internal orienting is not uniform but can change flexibly even within the same task context. The finding constrains theoretical models and motivates targeted investigations into the multiple processes that can support internal attention to guide adaptive behavior.

## Data Availability Statement

The code for this study is available on GitHub at https://github.com/Daniel-Gong/Dissociable-Neural-Signatures-WM-LTM. Data is available on Open Science Framework at https://osf.io/gk56s.

## Acknowledgements

This research was funded by a Clarendon Scholarship, a Medical Research Council Studentship, a New College-Yeotown Scholarship, and Yale PhD Scholarships to D.G.; a Wellcome Trust Senior Investigator Award (104571/Z/14/Z) and a James S. McDonnell Foundation Understanding Human Cognition Collaborative Award (220020448) to A.C.N. For the purpose of open access, the author has applied a CC BY public copyright licence to any Author Accepted Manuscript version arising from this submission.

## Author contributions

Dongyu Gong: Conceptualization; Investigation; Data curation; Formal analysis; Visualization; Project administration; Writing—Original draft; Writing—Review & editing. Dejan Draschkow: Conceptualization; Project administration; Resources; Supervision; Writing—Review & editing. Anna C. Nobre: Conceptualization; Funding acquisition; Project administration; Resources; Supervision; Writing—Original draft; Writing—Review & editing.

## Supplemental Information

**Figure S1.**
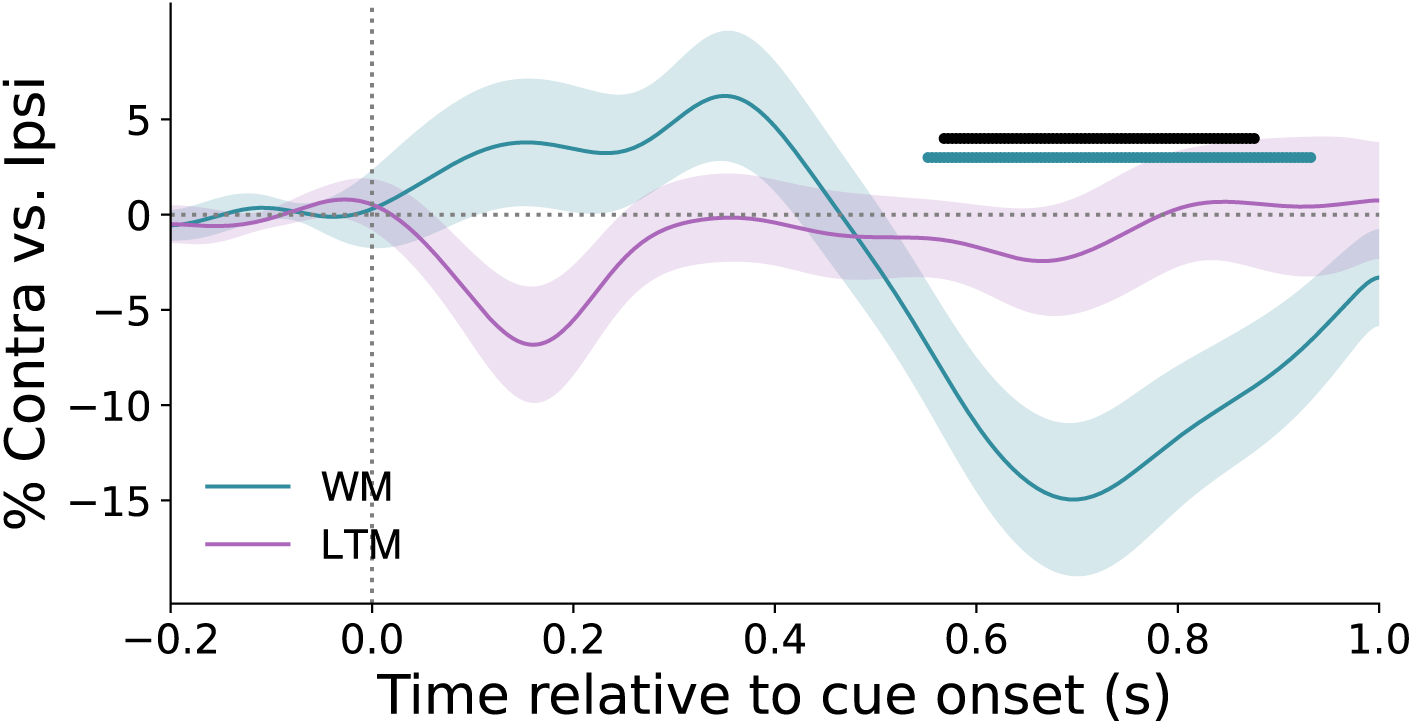
Time courses of alpha lateralization in the alpha band (8-12 Hz). Same analysis as in Fig. 3c but on EOG-regressed EEG data. Horizontal lines represent significant clusters (teal blue: WM compared to 0; black: WM vs. LTM difference). Shading areas represent ±1 SEM.

**Figure S2.**
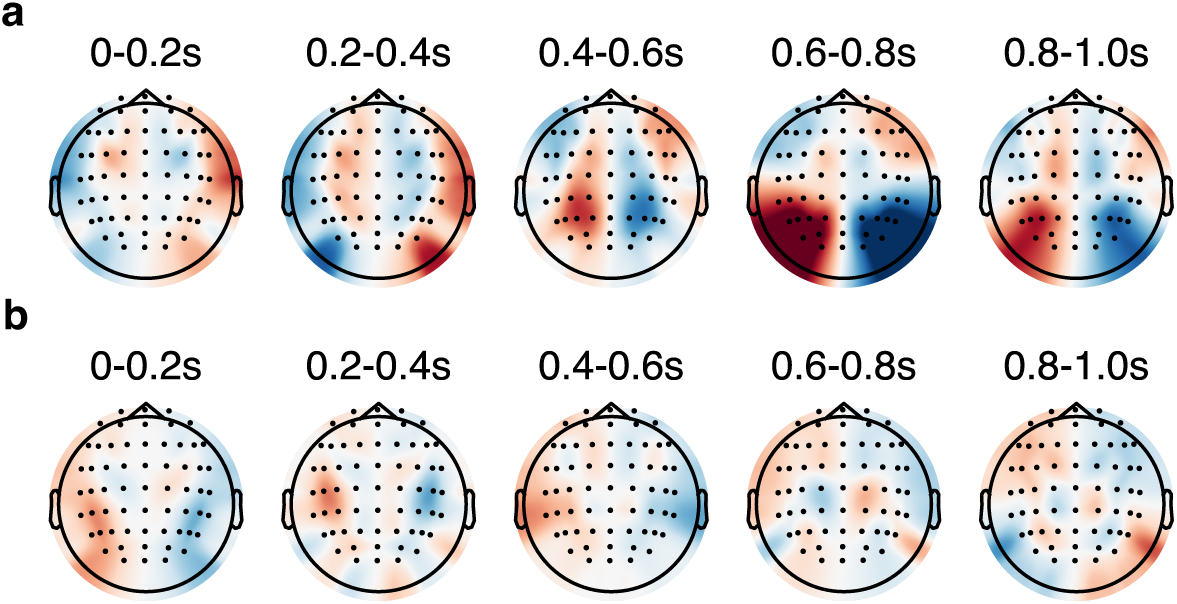
(a) Topographies of alpha power difference contralateral vs. ipsilateral to the memorized location in PO7/PO8 for cued WM items. The two halves of the topography are symmetrical, representing ipsilateral minus contralateral (left hemisphere) and contralateral minus ipsilateral (right hemisphere), respectively. (b) Same as (a) but for LTM items.

**Figure S3.**
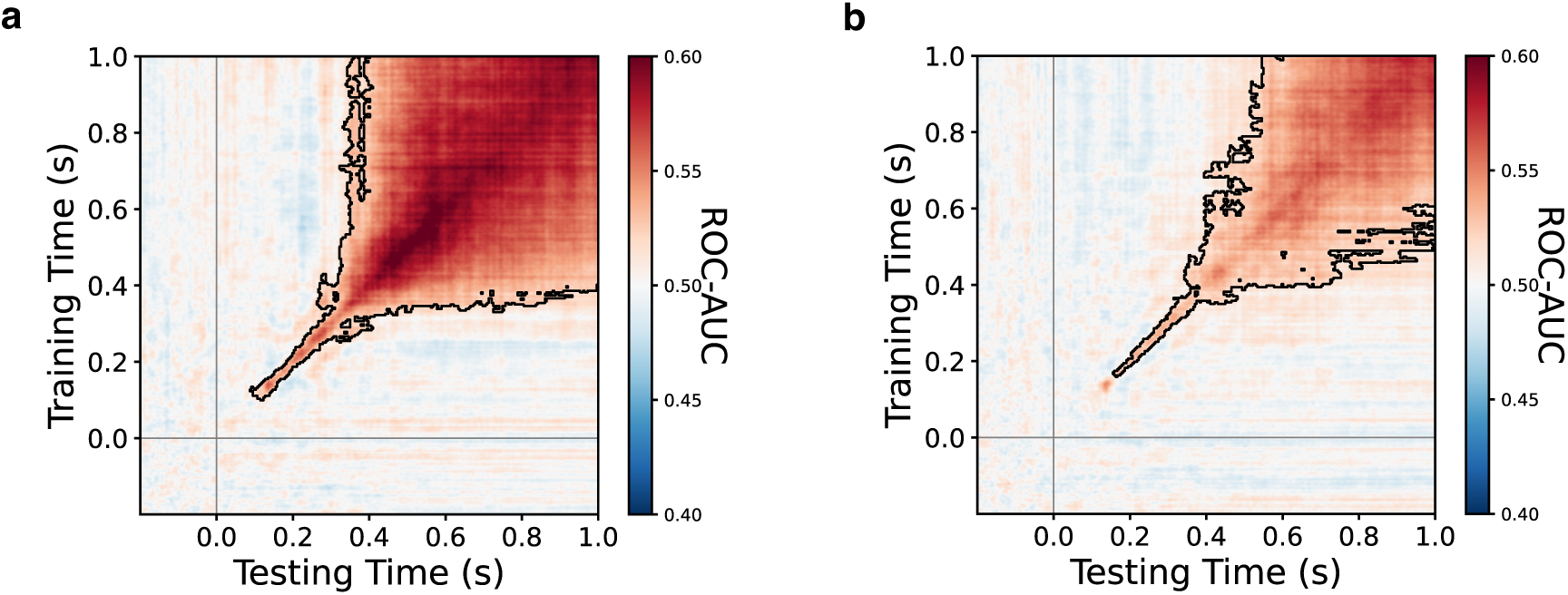
Temporal generalization matrix of classifier performance in discriminating between (a) neutral vs. WM and (b) neutral vs. LTM conditions. The diagonals correspond to the time course in Figure 4a. Black outlines in the matrix indicate significant clusters of temporal generalization above chance level.

**Figure S4.**
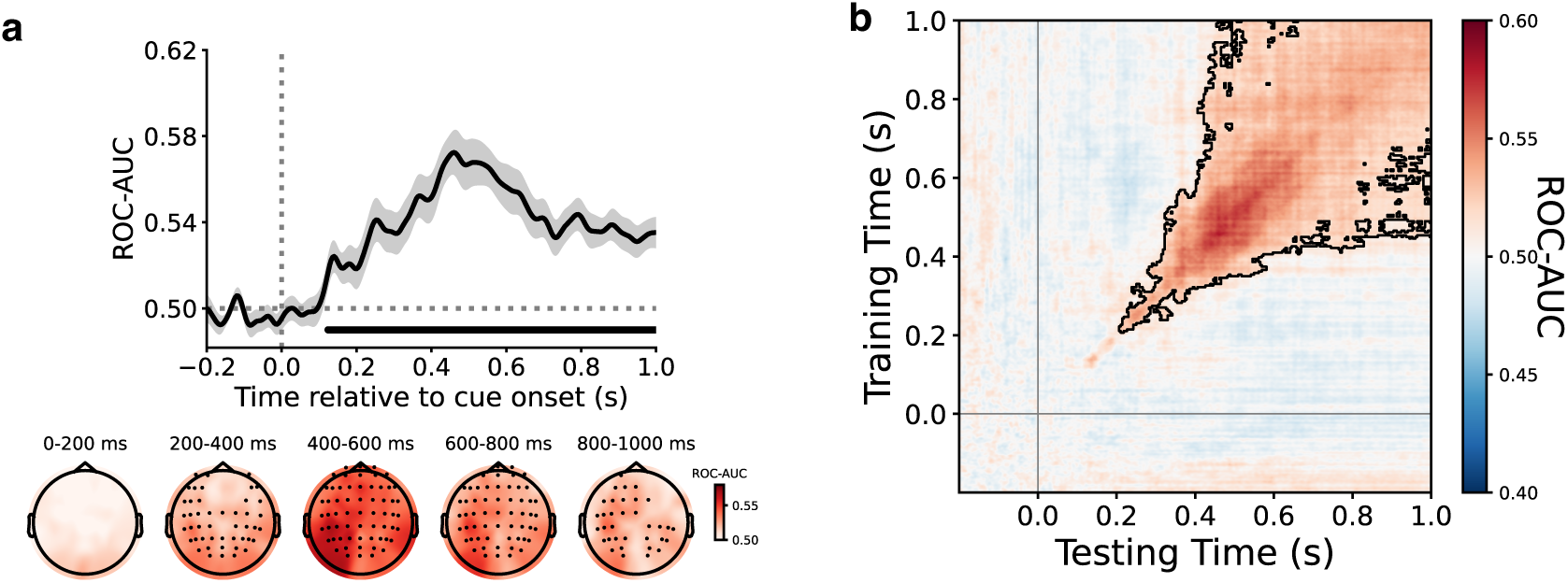
(a) To compare neural dynamics of selecting WM and LTM contents in a more straightforward way, we also trained a classifier to distinguish WM trials from LTM trials directly. As expected, we found reliable decoding of WM-vs. LTM-item conditions (cluster *p* = 0.001). The decoding topographies showed a widespread distribution of sensors that contributed significantly to the classification performance. (b) Temporal generalization matrix of classifier performance in discriminating between WM and LTM conditions. The diagonal corresponds to the time course in (a). Black outlines in the matrix indicate significant clusters of temporal generalization above chance level. Significant clusters in the temporal generalization matrix mainly existed along the diagonal at first. However, the temporal spread increased around 400 ms after cue onset (cluster *p* = 0.002), implying that the neural activity patterns distinguishing WM and LTM cueing conditions were initially more dynamic but became more stable over time.

**Figure S5.**
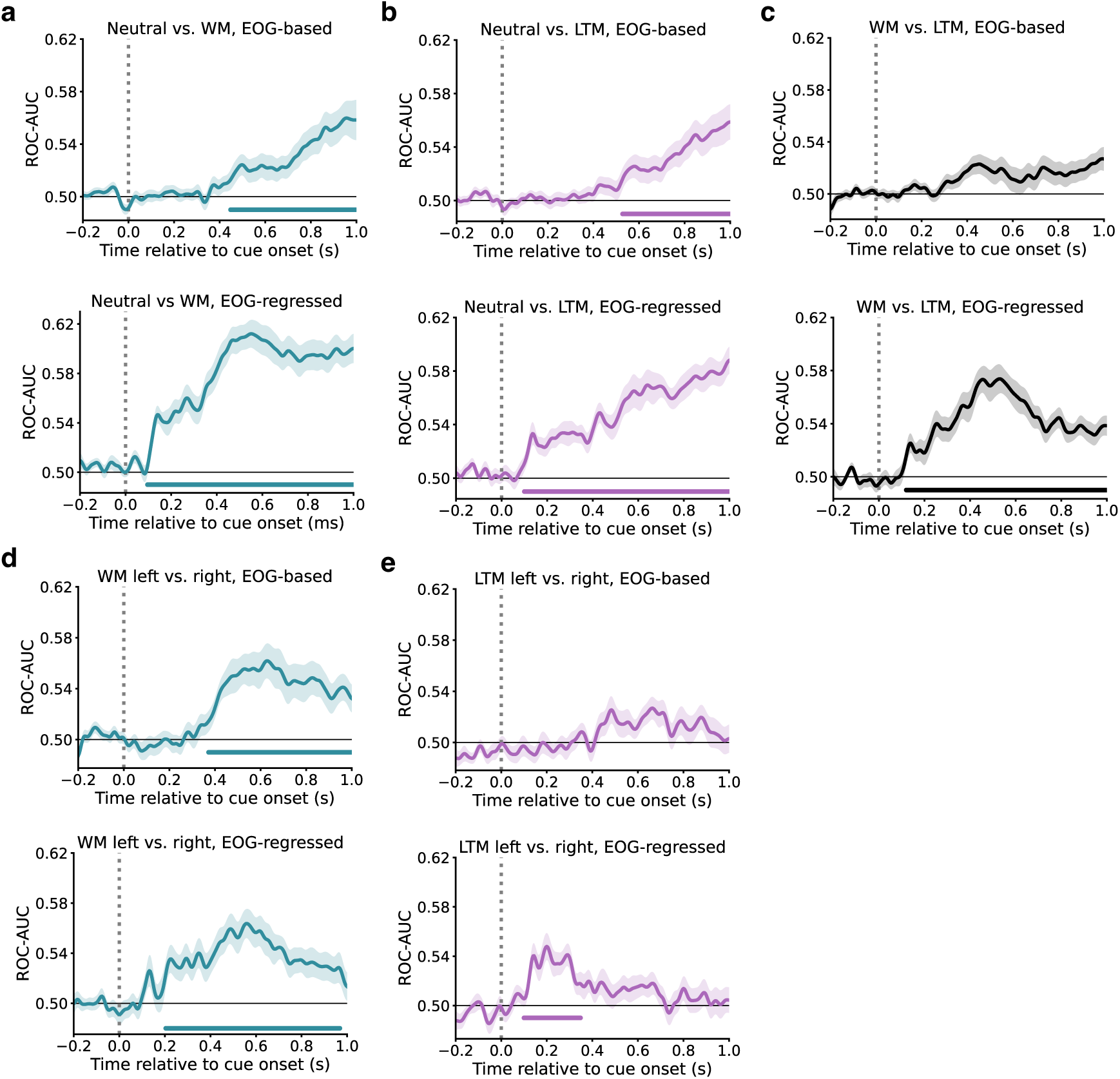
Control analysis examining the influence of eye movements on decoding scores. (a)-(e) show EOG-based decoding and EOG-regressed decoding scores for neutral vs. WM, neutral vs. LTM, WM vs. LTM, WM left vs. right, and LTM left vs. right, respectively. Shading areas represent ±1 SEM. Horizontal lines indicate significant temporal clusters.

**Figure S6.**
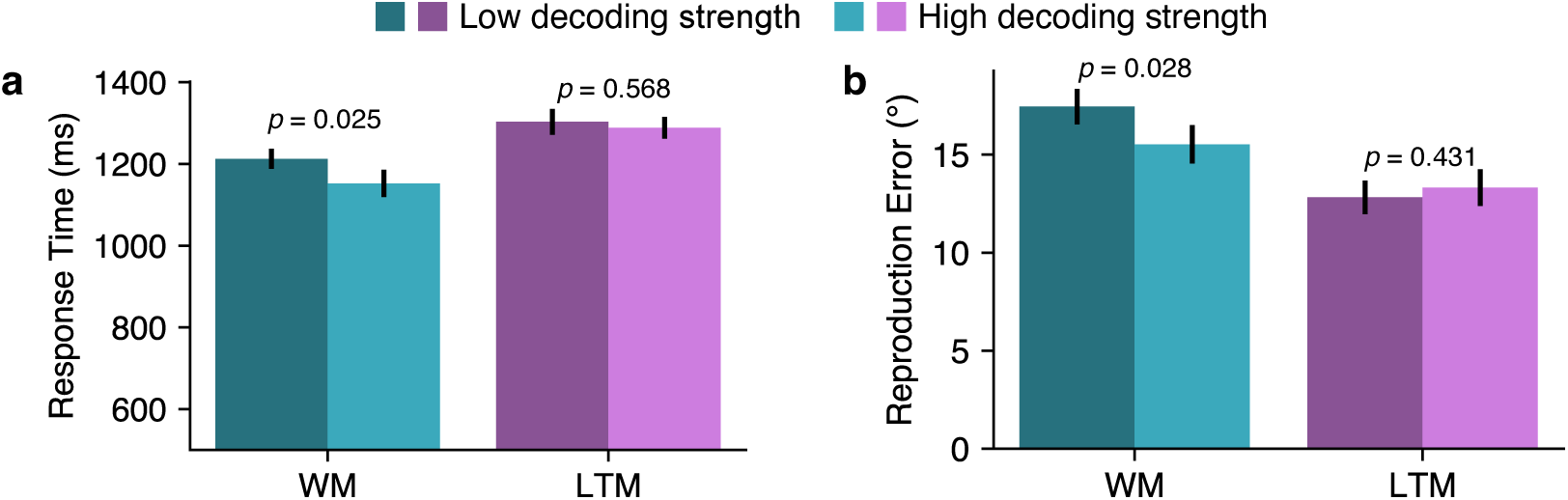
Relationship between decoding and behavioral performance. (a) WM and LTM response times as a function of trials with low vs. high decoding strength (against neutral) based on median split. (b) WM and LTM reproduction errors as a function of trials with low vs. high decoding strength (against neutral) based on median split.

